# Neural Representations of Post-Decision Choice Confidence and Reward Expectation in the Caudate Nucleus and Frontal Eye Field

**DOI:** 10.1101/2022.09.12.507621

**Authors:** Yunshu Fan, Takahiro Doi, Joshua I. Gold, Long Ding

## Abstract

Performance monitoring that supports ongoing behavioral adjustments is often examined in the context of either choice confidence for perceptual decisions (i.e., “did I get it right?”) or reward expectation for reward-based decisions (i.e., “what reward will I receive?”). However, our understanding of how the brain encodes these distinct evaluative signals remains limited because they are easily conflated, particularly in commonly used two-alternative tasks with symmetric rewards for correct choices. Previously we reported behavioral and neural results related to decision formation by monkeys performing a visual motion discrimination task with asymmetric rewards (Doi et al., 2020; Fan et al., 2020). Here we leveraged this task design to partially decouple trial-by-trial estimates of choice confidence and reward expectation and examine their impacts on behavior and their representations in the caudate nucleus (part of the striatum in the basal ganglia) and the frontal eye field (FEF, in prefrontal cortex). We found that these evaluative signals had infrequent, but consistent, effects on the behavior of well-trained monkeys. We also identified distinguishable representations of the two types of signals in FEF and caudate neurons, including different distribution patterns, time courses, and relationships to behavior in the two brain areas. These results suggest that the cortico-striatal decision network may use diverse evaluative signals for performance monitoring and add to our understanding of the different roles of the FEF and caudate nucleus in decision-related computations.

## Introduction

Effective learning can depend on comparisons between expected and experienced outcomes (Sutton and Barto, 1998). These expectations represent a form of performance monitoring that often takes one of two forms: choice confidence or reward expectation. Confidence is the subjective belief that a choice is correct given the evidence (Kiani et al., 2014a) and is commonly used to evaluate decisions based on unreliable or noisy sensory evidence. It is also thought to support adaptive strategies in changing environments and account for other forms of sequential behavioral adjustments including post-error slowing (Yu and Dayan, 2005; Nassar et al., 2012; Purcell and Kiani, 2016). Reward expectation is the expected benefit (and/or cost) given a choice. It is a critical component of reinforcement learning and is commonly used to evaluate value-based decisions (Sutton and Barto, 1998; Samejima et al., 2005; Daw and Doya, 2006; Rangel et al., 2008; Schultz, 2015).

Neural signals consistent with one or the other of these two forms of evaluative signals for performance monitoring have been reported in many brain regions, including the caudate nucleus of the basal ganglia and the frontal cortex (for a limited sample, Kawagoe et al., 1998; Schultz, 1998; Roesch and Olson, 2003; Padoa-Schioppa and Assad, 2006; Kepecs et al., 2008a; Lau and Glimcher, 2008; Kiani and Shadlen, 2009a; Basten et al., 2010; Ding and Gold, 2010; Nomoto et al., 2010; Kennerley et al., 2011; Middlebrooks and Sommer, 2012; Teichert et al., 2014; Yanike and Ferrera, 2014a; Hebart et al., 2016; So and Stuphorn, 2016; Lak et al., 2017, 2020b). However, our understanding of these neural representations of confidence and reward expectation has been limited by the fact that these quantities are easily conflated under conditions in which they are typically examined. For example, for value-based decision tasks, choice confidence can be based on a comparison of reward expectations for the chosen versus the unchosen options. So and Stuphorn (2016) tackled this problem with dissociable risk and reward manipulations and clarified that neural activity in supplementary eye field encoded choice confidence instead of reward expectation. Likewise, for many perceptual-decision tasks, the reward expectation for the chosen option is the product of the confidence in making a correct choice and the magnitude of reward associated with a correct choice. When the reward magnitude is fixed, choice confidence and reward expectation are perfectly correlated. Because of these potential confounds, if and how choice confidence and reward expectation have distinct representations in the brain, including in caudate and FEF, is not well understood in the context of perceptual decision-making.

**Figure 1.**
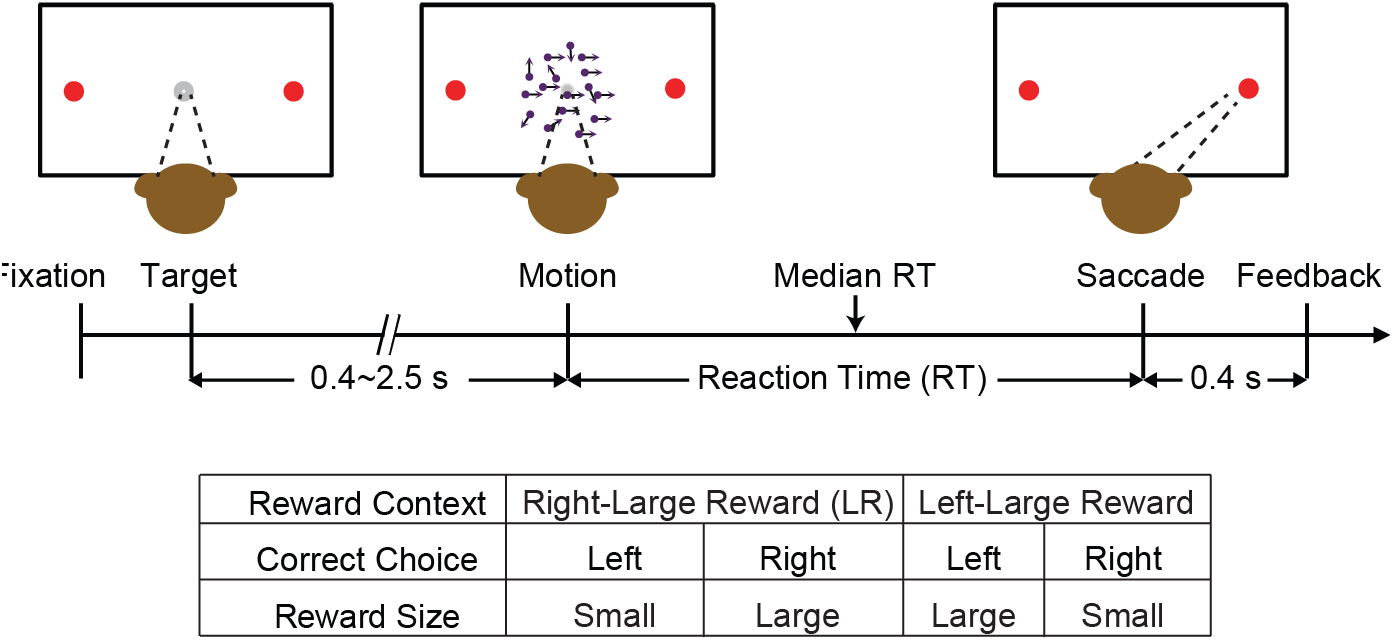
Task design and timeline. Monkeys reported the perceived motion direction with a saccade to one of the two choice targets. The motion stimulus was turned off upon detection of saccade. Correct trials were rewarded based on the reward context. Error trials were not rewarded.

To address this issue, we leveraged a behavioral task with separate manipulations of evidence strength and reward-choice associations (Figure 1A). As we show below, this task design uncoupled the estimated choice confidence and reward expectation, thus allowing us to differentiate neural representations of the two quantities at the single-neuron level in the caudate and FEF. We previously showed that neurons in these two areas play similar, but distinguishable, computational roles in forming these decisions that require balancing uncertain sensory evidence with asymmetric reward expectations (Fan et al., 2020). Here we show that these regions may also play similar, but distinguishable, roles in monitoring and adjusting these decisions, by keeping track of both choice confidence and reward expectation.

## Results

### Reward asymmetry-induced bias can help distinguish between confidence and reward expectation

For a typical perceptual-decision task, choice confidence is estimated as the expected accuracy as a function of choice and decision time (Kiani and Shadlen, 2009a; Fetsch et al., 2014; Kiani et al., 2014a). Reward expectation is estimated as the product of confidence and reward magnitude associated with the chosen option. When the reward magnitude is equal between the two choices, choice confidence and reward expectation are perfectly correlated. To illustrate this relationship, we simulated a decision process with the drift diffusion model (DDM) that makes decisions based on the accumulation of motion evidence over time. As shown previously, both quantities are higher for choices made with shorter decision times and when evidence is stronger (higher coherence; Figure 2A-E, top row, black lines; Kepecs et al., 2008a; Kiani and Shadlen, 2009a; Kiani et al., 2014b; Lak et al., 2019).

**Figure 2.**
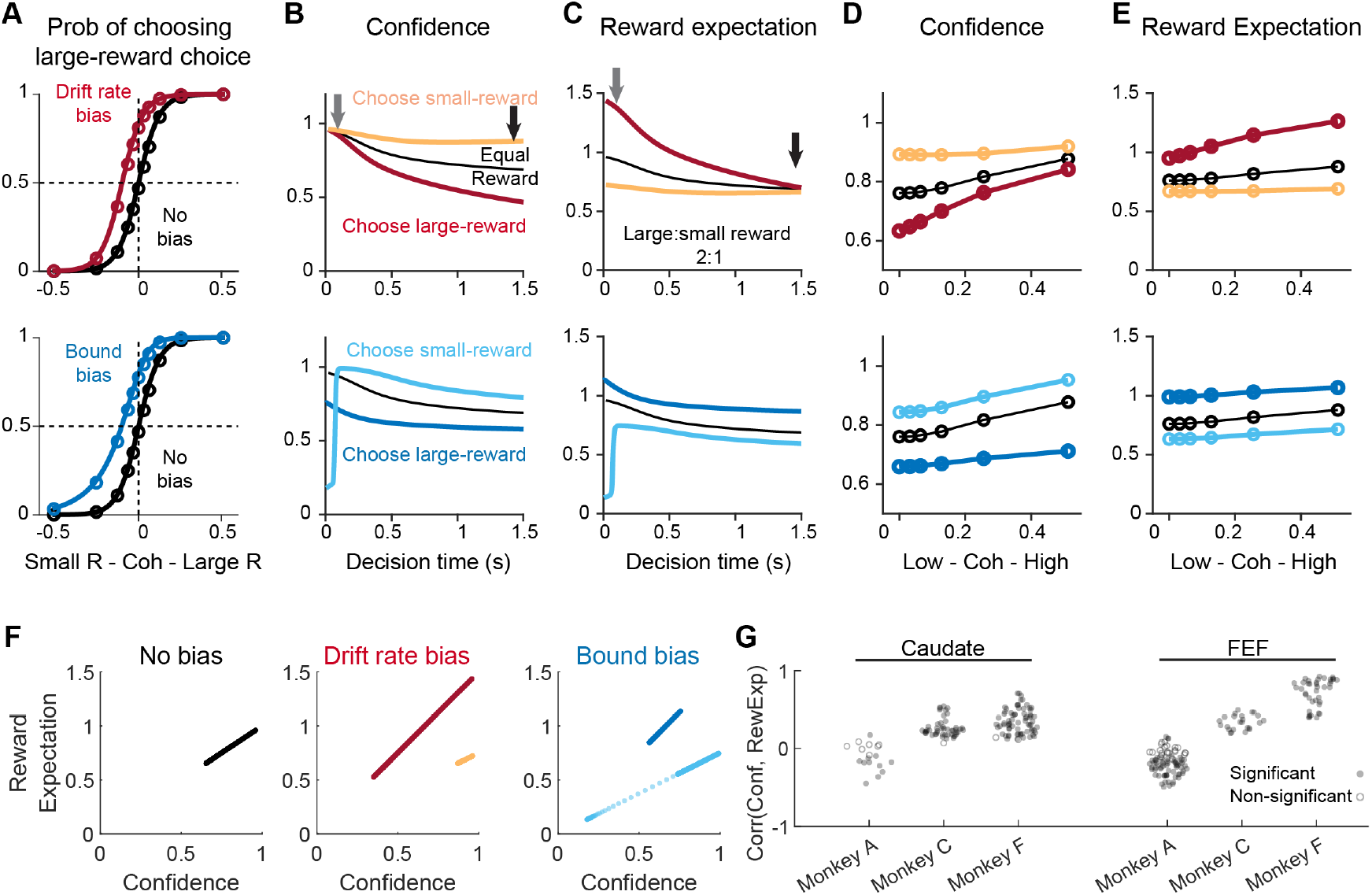
Confidence and reward expectation depended on decision time, motion coherence, and reward asymmetry-induced biases. **(A-F)** DDM simulation results for three scenarios: 1) no bias with equal reward for both choices (black), 2) a drift rate bias favoring the large-reward option (red/orange, top row and F), and 3) a bound bias favoring the large-reward option (blue, middle row and F). DDM parameters: *total bound height* = 2; *scaling factor* = 8; *drift-rate bias* (top) = 0.1; *bound bias* (i.e., the ratio between the large-reward bound and small-reward bound; bottom) = 4. Drift-rate bias and bound bias values were chosen to generate similar horizontal shifts of the psychometric function given ambiguous evidence (coherence = 0). **(A)** Probability of a large-reward choice as a function of signed motion coherence. Negative/positive coherence: motion towards the small-/large-reward choice. Circles indicate the coherence levels used in the simulation. **(B and C)** Choice confidence and reward expectation as a function of decision time, pooled across all coherences. **(D and E)** Choice confidence and reward expectation as a function of coherence, pooled across all decision times. **(F)** Scatterplots of relationships between choice confidence and reward expectation given no bias (left), a drift-rate bias (middle), and a bound bias (right). **(G)** Distribution of Pearson correlation coefficients between confidence and reward expectation in each recording sessions from the three monkeys and two brain regions. Note that all data points are < 1 (where 1 = perfect correlation for the equal-reward condition).

When the two choices are associated with different reward magnitudes, a subject should use a bias towards the larger-reward choice (particularly when the sensory evidence is weak) to maximize overall reward rate. In the DDM framework, such a choice bias is modeled as a bias in evidence accumulation (drift-rate bias), decision bounds (bound bias), or both. The two forms of bias affect choice confidence and reward expectation functions differently and in a choice-dependent manner (Figure 2A–E, compare the top and middle rows). For example, with a drift-rate bias favoring the large-reward choice and compared to the equal-reward condition, choice confidence is overall higher, less dependent on decision time, and less dependent on evidence strength for the smallreward choice (Figure 2B and D, orange curves are higher and flatter than black curves). Choice confidence is overall lower, more dependent on decision time, and more dependent on evidence strength for the large-reward choice (Figure 2B and D, red). The higher overall confidence for small-reward choices reflects the tendency that these choices are made only when the large-reward choice is more likely to be incorrect. With a bound bias, choice confidence for the small-reward choice is much lower for very fast choices, higher for other decision times and higher across evidence strength (Figure 2B, light blue). Choice confidence for the large-reward choice is lower overall and less dependent on decision time and evidence strength (Figure 2B, dark blue).

More importantly, regardless of which bias is in play, with asymmetric rewards choice confidence and reward expectation no longer follow identical relationships with decision time and evidence strength (compare Figure 2B and C, D and E). For example, with a moderate large-to-small reward ratio of 2:1 and a drift-rate bias, fast decisions with almost identical choice confidence can have substantially different reward expectations for the small- and large-reward choices (Figure 2B and C, gray arrows). Moreover, slow decisions with very different choice confidence can have similar reward expectations (black arrows). For both types of biases, small-reward choices tend to show higher choice confidence and lower reward expectation, and vice versa for large-reward choices. The de-coupling between choice confidence and reward expectation can also be summarized with scatterplots of the two quantities (Figure 2F). In contrast to the perfect correlation for the unbiased DDM in the equal-reward condition, the two quantities are perfectly correlated only within a reward type and not across reward types. We therefore used comparisons across reward types to distinguish the behavioral and neural sensitivity to choice confidence and reward expectation.

### Monkeys showed partially decoupled choice confidence and reward expectation

We trained three monkeys to perform a response-time (RT), asymmetric-reward, random-dot visual motion direction-discrimination saccade task (Figure 1). The monkeys were presented with a random-dot kinematogram and required to make saccades indicating their judgements about the global motion direction. Across trials, two directions (left or right) and five motion strengths were interleaved pseudo-randomly. We manipulated the choice-reward associations (reward context) in a block design. In “Right-Large Reward (LR)” blocks (~50 trials per block), a correct rightward saccade was paired with a large reward, and a correct leftward saccade was paired with a small reward. In “Left-LR” blocks, a correct leftward saccade was paired with a large reward, and a correct rightward saccade was paired with a small reward. Error trials were not rewarded. The two types of blocks were alternated in a session, and the identity of the upcoming block was signaled to the monkey at each block change. As we documented previously, the three monkeys showed consistent behavioral strategies such that their choice and response time (RT) depended on both the reward context and motion strength, and their reward-biased decision strategy can be captured with a combination of drift-rate and bound biases in DDM model fits (Fan et al., 2018; Doi et al., 2020).

We analyzed behavioral and neural data from 132 sessions with caudate recordings (*n*=17, 45, and 70 from monkey A, C, and F, respectively) and, separately, 131 sessions with FEF recordings (*n*=75, 23, and 33 from monkey A, C, and F, respectively). We estimated choice confidence and reward expectation for each trial using DDM fits to each monkey’s session-by-session choice and RT data. For each individual session, these estimates of choice confidence and reward expectation were not perfectly correlated with each other and therefore at least partly distinguishable, with monkey A showing the weakest correlation and monkey F showing the strongest correlation (Figure 2G).

### Influence of confidence and reward expectation on subsequent decisions in well-trained monkeys

All three monkeys were well trained on the task and made choices whose accuracy and speed could be well accounted-for via the DDM; i.e., they were based primarily on a decision process that combined the accumulated sensory evidence on the current trial with certain reward context-dependent biases (Fan et al., 2018; Doi et al., 2020). Nevertheless, the monkeys also exhibited some sequential (trial-to-trial) behavioral effects that reflected choice confidence, reward expectation, or both. We assessed these sequential effects using: 1) logistic regression testing for effects on staying or switching on the subsequent choice, and 2) linear regression testing for effects on speeding up or slowing down the subsequent decision process.

The monkeys exhibited sequential effects that depended on choice confidence in 30 out of 263 sessions, with consistent patterns (Figure 3A-C). When the previous correct choice was associated with higher choice confidence, the monkeys were more likely to repeat that choice (Figure 3A, Wilcoxon signed-rank test for *H_0_*: the median value of the regression coefficient=0, *p*=0.014) and speed up the next choice (Figure 3B; *p*<0.0001). There was no additional modulation on RT when the same choice was repeated (Figure 3C; *p*>0.5). Likewise, the monkeys exhibited sequential effects that depended on reward expectation in 41 out of 263 sessions, with similarly consistent patterns: the monkeys were more likely to repeat a correct choice associated with high reward expectation and speed up the next choice (Figure 3D–F, *p*<0.0001, =0.027, and >0.5, respectively).

**Figure 3.**
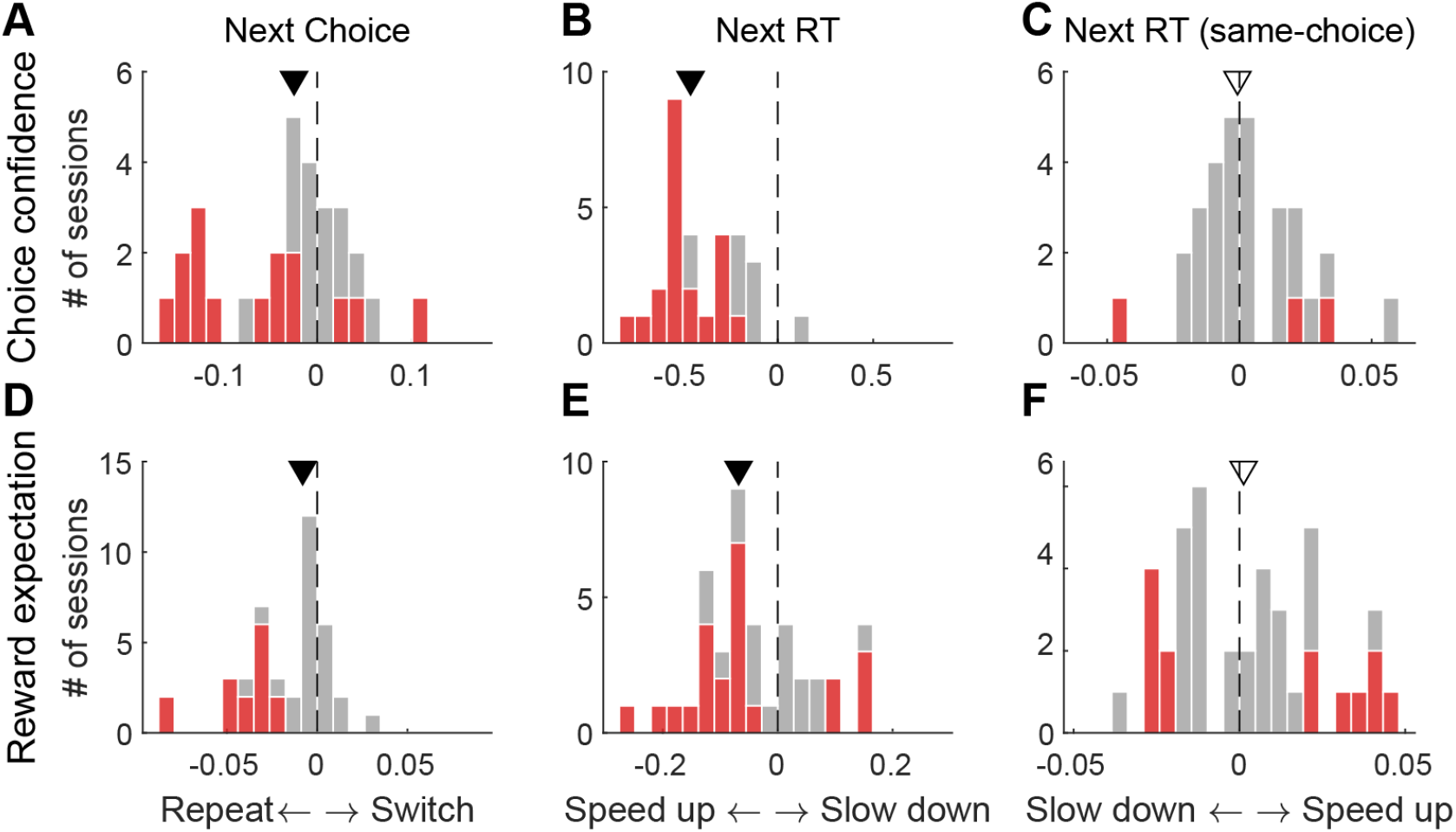
Sequential effects associated with choice confidence and reward expectation. **(A–C)** Histograms of regression coefficients for sequential effects on the next choice (A; β_*prevConf*_), RT independent of choice (B; α_*prevconf*_), and RT of only repeated choices (C; α_*prevConf×Switch*_). Only sessions with significant sequential effects associated with choice confidence were included (*n* = 30). Triangle: median value; filled: *p*<0.05, Wilcoxon signed-rank test. Red bars: sessions with significant non-zero coefficients; gray bars: sessions with coefficient not significantly different from zero. **(D–F)** Same format as A–C, for sequential effects associated with reward expectation (*n* = 41).

We did not find any temporal (across-session) clustering of these effects (Figure 3-Supplement 1). Monkey F appeared more likely to rely on reward expectation than choice confidence, whereas the other two monkeys had mixed dependences. These results indicate that even in well-trained monkeys, choice confidence and reward expectation can occasionally have measurable behavioral effects that include trial-by-trial adjustments of choice and RT.

### Choice confidence and reward expectation are reflected in post-decision activities of FEF and caudate neurons

We previously showed that in monkeys performing this task, caudate and FEF neurons encode combinations of motion coherence, decision time, and reward size and context in time periods just before and after the saccadic response; i.e., after the decision has been made (Doi et al., 2020; Fan et al., 2020). Here we show that these neurons also encode post-decision evaluative signals including choice confidence and reward expectation that depend on particular combinations of these factors (Figure 2).

Figures 4 and 5 show example caudate and FEF neurons, respectively, with peri-saccade activity that correlates with choice confidence or reward expectation. The caudate neuron in Figure 4A had activity that was the highest around the time of saccade to the ipsilateral choice target and was modulated by motion coherence (shade) and the expected reward size (color). Its average firing rate for the preferred choice increased with motion coherence, decreased with decision time, and was higher for the small-reward choice, consistent with a confidence signal. The example caudate neuron in Figure 4B was not as choice selective as the neuron in Figure 4A. Its firing rate for the preferred choice decreased with motion coherence, increased with decision time, and was higher for the large-reward choice, consistent with a negative confidence signal. The example caudate neuron in Figure 4C had pre-saccadic activity that increased with motion coherence, decreased with decision time, and was higher for the large-reward choice, consistent with a reward expectation signal. The example caudate neuron in Figure 4D had post-saccade activity with opposite modulation patterns, consistent with a negative reward expectation signal. Single-neuron examples with similar modulation patterns in FEF are shown in Figure 5.

**Figure 4.**
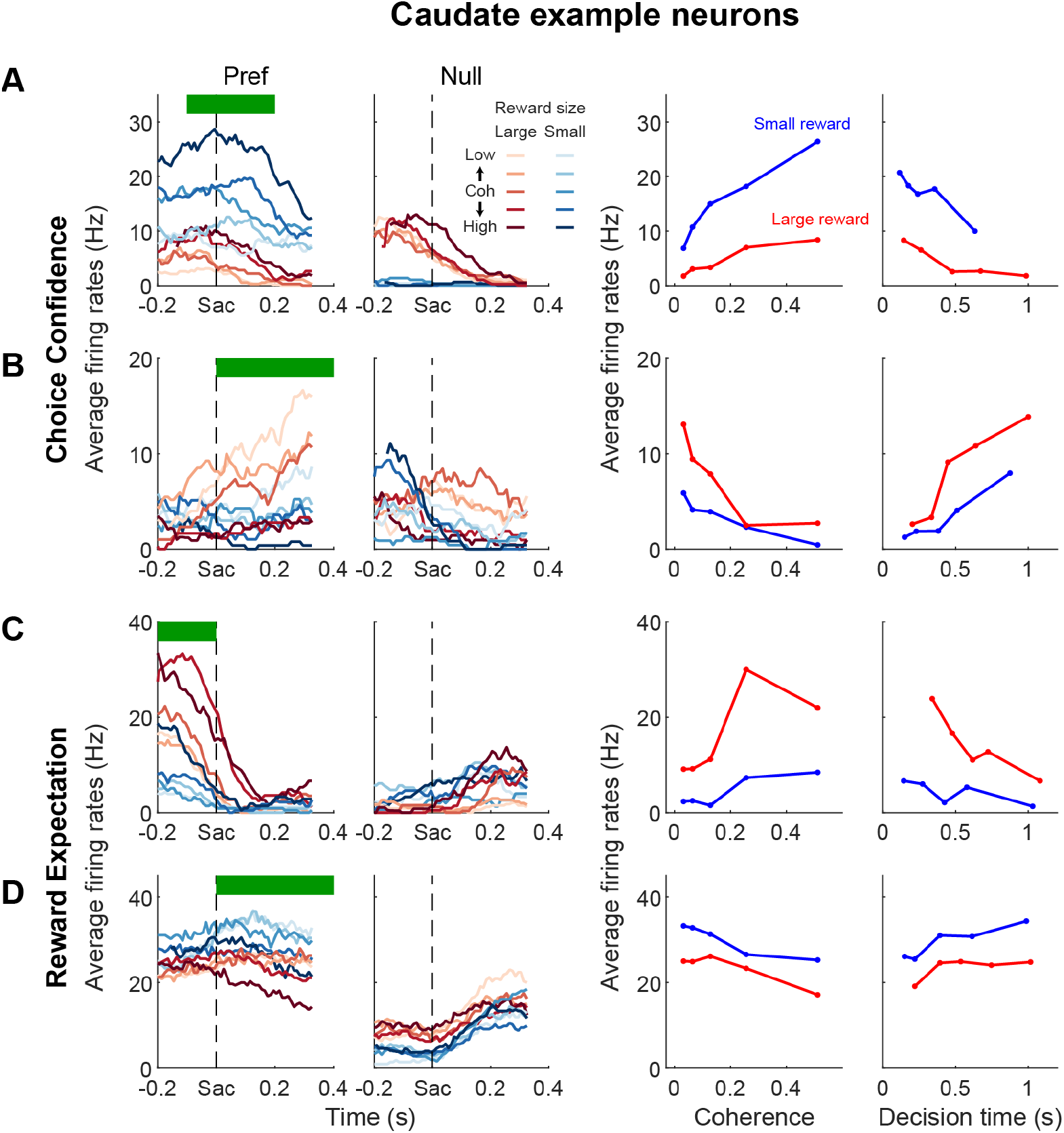
Example caudate neurons encoding confidence or reward expectation. Each row shows the activity of one neuron. Columns 1 and 2 plot the average firing rates for its preferred and null choices for different combinations of coherence and expected reward size. Shades: coherence levels. Colors: reward size. Firing rates were computed for correct trials only, using a 200ms running window (50-ms steps). Columns 3 and 4 plot the average firing rates in the epoch defined by the green bar in the first column as a function of coherence and decision time, respectively. Decision time was obtained from DDM fits to session-specific choice and RT behavior and grouped into five quantiles.

**Figure 5.**
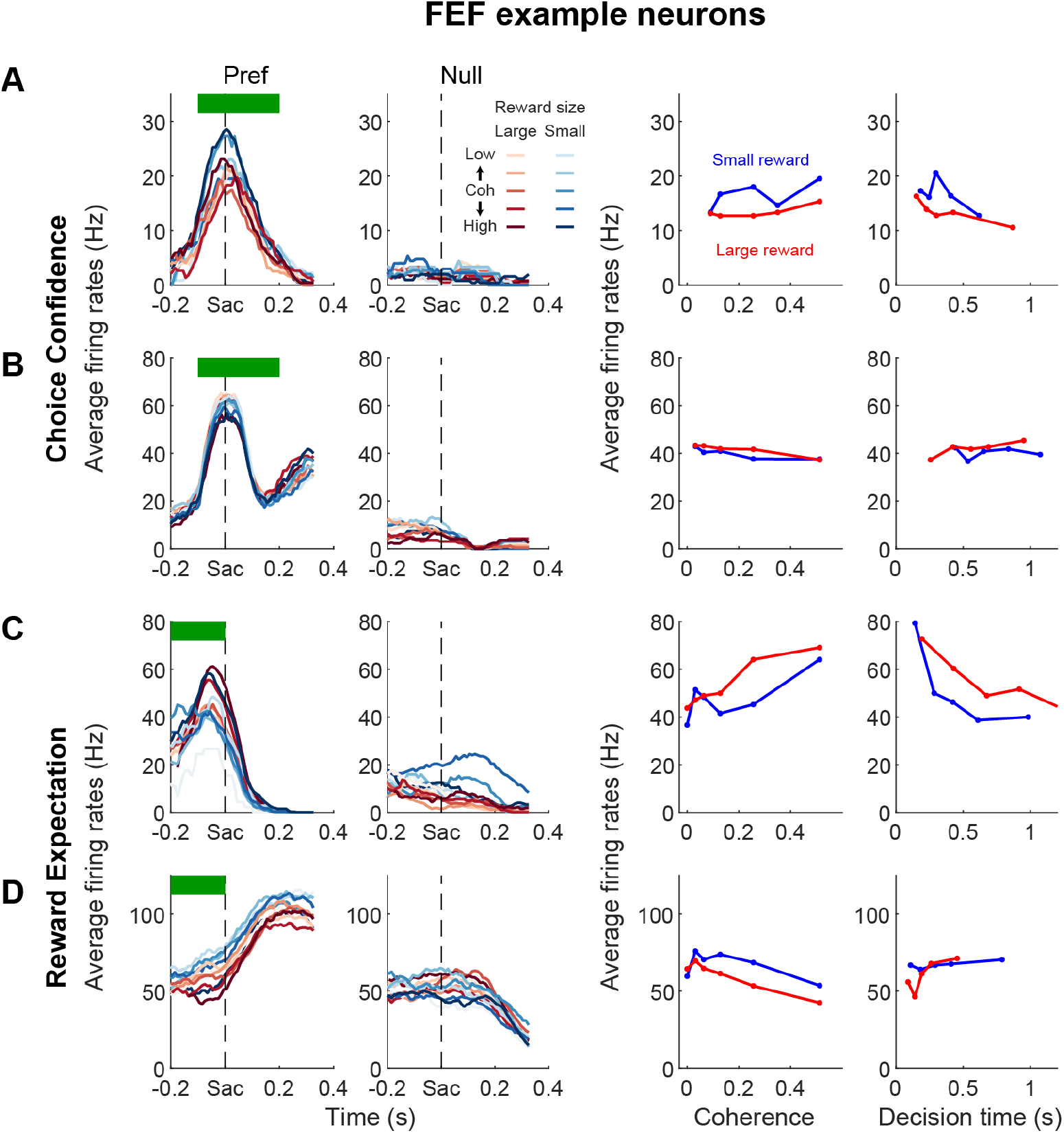
Example FEF neurons encoding confidence or reward expectation. Same format as Figure 4.

To test whether a neuron’s activity was sensitive to choice confidence or reward expectation, we computed two partial correlations between firing rate and each measure, while accounting for the other. We performed this analysis for each choice separately. We observed significant non-zero partial correlation coefficients for choice confidence or reward expectation in many neurons in both caudate and FEF samples. Some of these neurons showed reliable choice selectivity in their activity around saccade onset, as tested previously using multiple linear regression (Fan, 2020; 100 ms before saccade onset to 200 after), whereas others did not. The heatmaps of correlation coefficients for the choice-selective and not-selective subpopulations are shown in Figure 6 and Figure 6-Supplement 1A, respectively.

**Figure 6.**
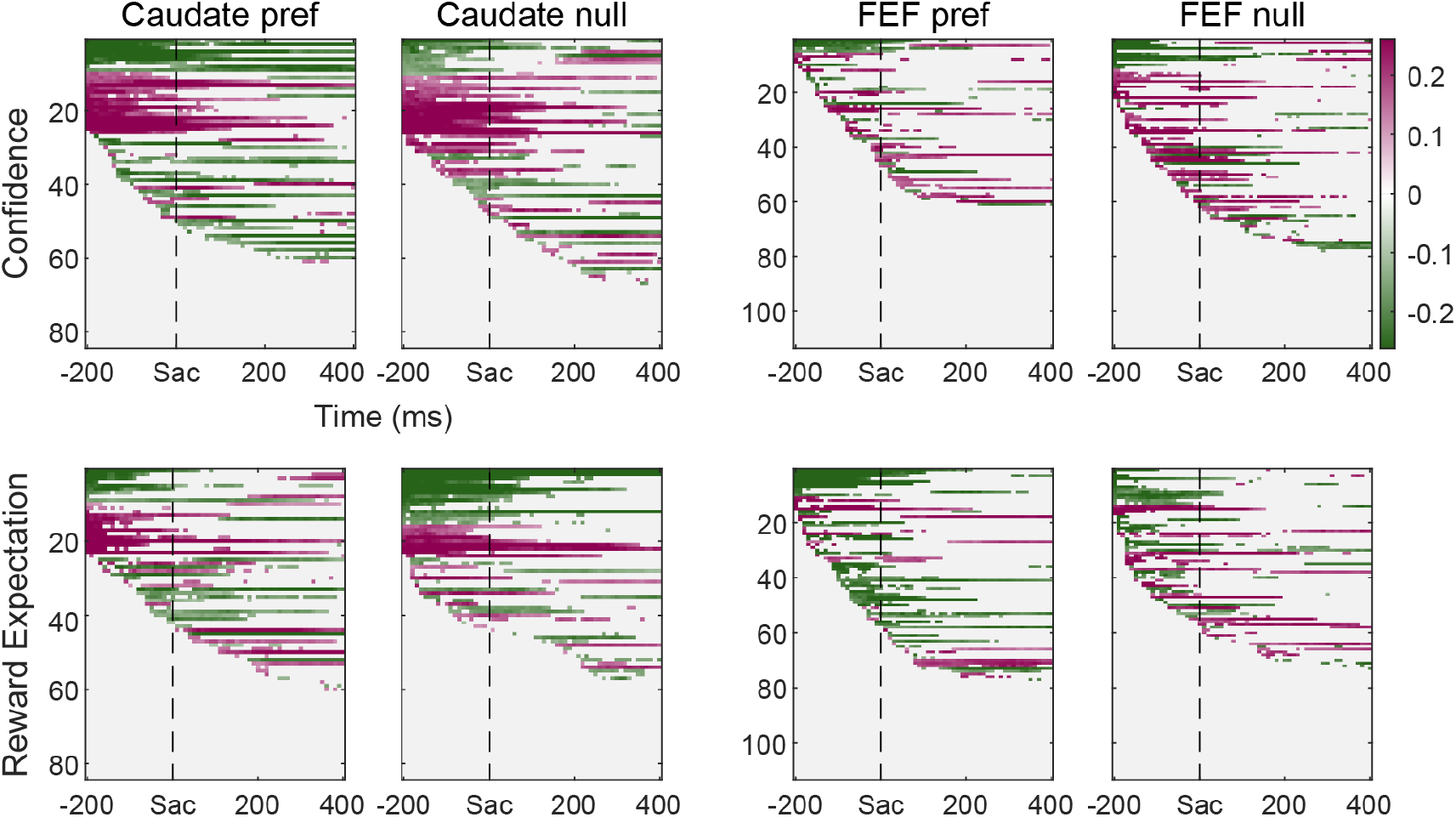
Partial correlation coefficients for neurons with choice-selective activity around saccade onset. Top: correlation coefficients between firing rates and choice confidence, after accounting for the effect of reward expectation. Bottom: correlation coefficients between firing rates and reward expectation, after accounting for the effect of choice confidence. Neurons are sorted by the onset of the significant nonzero coefficient and sign of the coefficient, separately for the preferred and null choices. Each pixel shows the result for average firing rates computed in a 300-ms running window (10-ms step).

Within each brain region and subpopulation, we did not observe significant differences in the prevalence of neural representations of choice confidence or reward expectation between trials with the preferred and null choices for the choice-selective neurons (Figure 7A,B, first two panels; Chi-square test, *p* > 0.05), nor between trials with contra- and ipsi-lateral choices for the choice non-selective neurons (Figure 6-Supplement 1B,C, first two panels). Choice confidence and reward expectation were also represented with similar prevalence in the two subpopulations of each brain region (Figure 7A,B and Figure 6-Supplement 1B, C, third panels).

**Figure 7.**
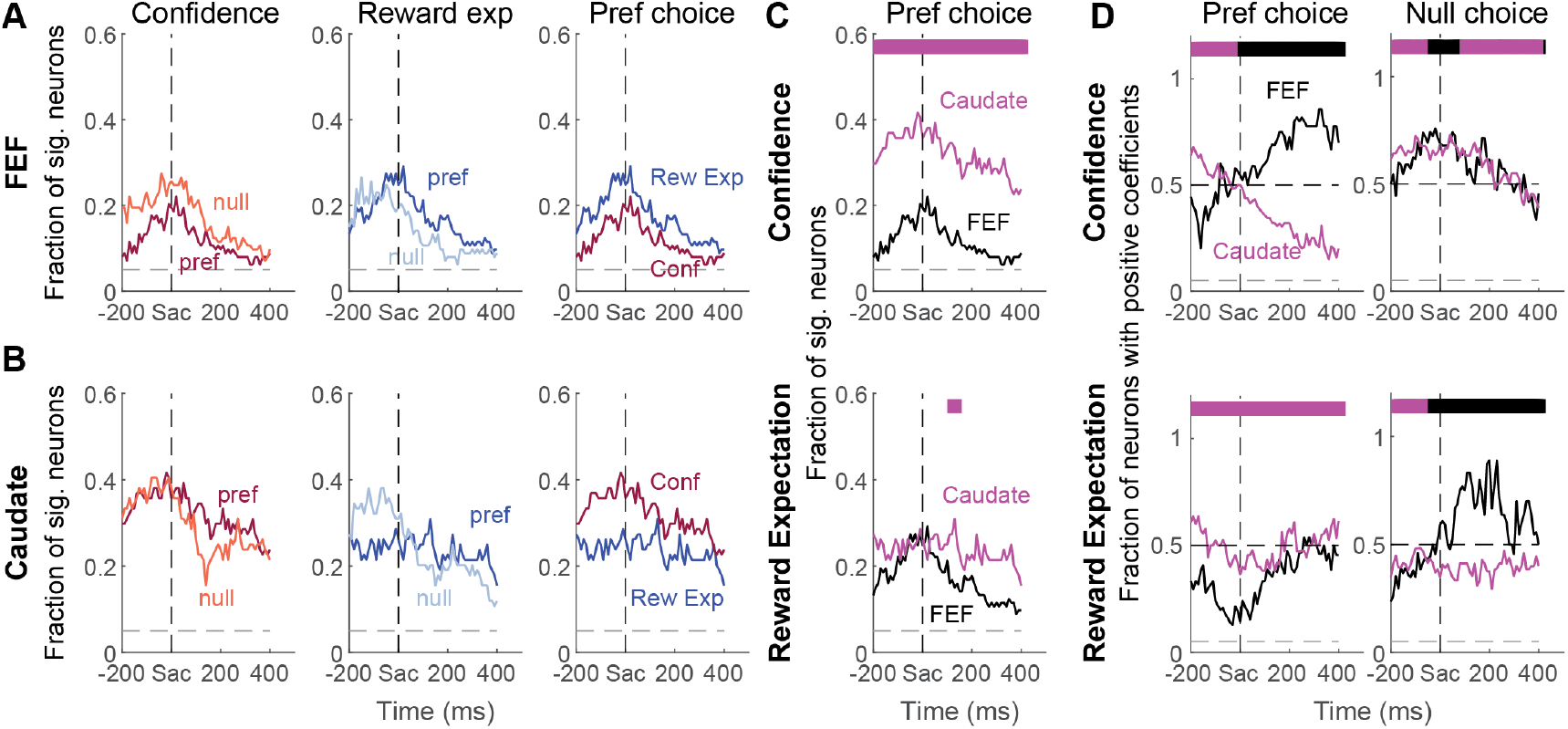
Fractions of significant non-zero partial correlation coefficients for neurons with choice-selective activity around saccade onset. **(A-B)** Comparisons of the prevalence of modulation between the preferred and null choice trials and between confidence and reward expectation for the preferred choice, in the FEF (A) and caudate samples (B), respectively. Dashed lines indicate chance level. **(C)** Comparisons of the prevalence of modulation by confidence (top) and reward expectation (bottom) between FEF and caudate samples. The bar on top of the curves shows the time points (in 12 bins) with significant differences between the two samples (color indicates the region with the larger fraction; Chi-square test *p*<0.05/12). **(D)** Comparisons of the prevalence of positive coefficients between FEF and caudate samples for the preferred and null choices separately. Same format as C. Only neurons with significant non-zero coefficients were included and time bins with fewer than six of such neurons were excluded.

Between caudate nucleus and FEF, we observed two regional differences. First, there was a higher prevalence of choice-confidence representation in the caudate sample than in the FEF sample (Figure 7C and S6D, top panels). Reward expectation representation was also more prevalent in the caudate sample, especially after saccade onset (bottom panels). Second, the dominant signs of correlation coefficients differed between the two regions. For neurons with choice-selective peri-saccade activity, the coefficients for choice confidence were primarily negative before saccade onset and positive afterward for FEF (Figure 7D). The opposite time course was observed for caudate neurons. Caudate neurons without choice-selective peri-saccade activity also showed a positive-to-negative dominant transition, with the transition occurring later than for caudate neurons with choice-selective activity (Figure 6-Supplement 1E). The coefficients for reward expectation were primarily negative before and around saccade onset for FEF neurons, whereas the signs were more evenly distributed throughout for caudate neurons. Together, these results suggest that evaluative signals were present in both FEF and caudate activity, but how these signals were represented differed between the two regions.

To assess the robustness of these results, we considered two potential caveats in the analyses. First, because choice confidence and reward expectation both depended on reward context, it is possible that neurons that encode reward context alone could appear to show a reliable correlation coefficient for either choice confidence or reward expectation. To counter this possibility, we imposed an additional criterion that neurons encoding evaluative signals must be also sensitive to decision time. This criterion filtered out a very small number of neurons and did not change the above-mentioned results qualitatively (Figure 7-Supplement 1). Second, our estimates of reward expectation were based on the ratio of delivered juice volume, which may differ from the monkeys’ subjective estimate of reward ratio. As an alternative approach, we defined the monkey’s internal estimate of reward ratio as the value that would lead to the largest correlation coefficient between the corresponding reward expectation and neural activity. We applied this approach to activity in three periods: pre-saccade, peri-saccade, and post-saccade. As shown in Figure 7-Supplement 2A and B, the “best” reward ratio tended to be near 1 for activity modulated primarily by choice confidence and could deviate substantially from the reward ratio in juice volumes. Using the best reward ratios, we repeated the partial correlation analysis and found that the regional differences in the prevalence of representations of choice confidence and reward expectation remained largely intact (Figure 7-Supplement 2C).

Lastly, we examined how the neural representations of evaluative signals related to how these signals were used for behavioral adjustments. Specifically, we computed partial correlation coefficients between: 1) neural activity and choice confidence after accounting for reward expectation (summing the measure in Figure 6 across both choices), for neurons recorded in sessions with choice confidence-dependent sequential effects on choice or RT (*n* = 9 and 25 neurons in caudate and FEF, respectively; heatmaps in Figure 8A and B); and 2) neural activity and reward expectation after accounting for choice confidence, for neurons recorded in sessions with reward expectation-dependent sequential effects on choice or RT (*n* = 27 and 18 neurons in caudate and FEF, respectively; Figure 8E and F).

**Figure 8.**
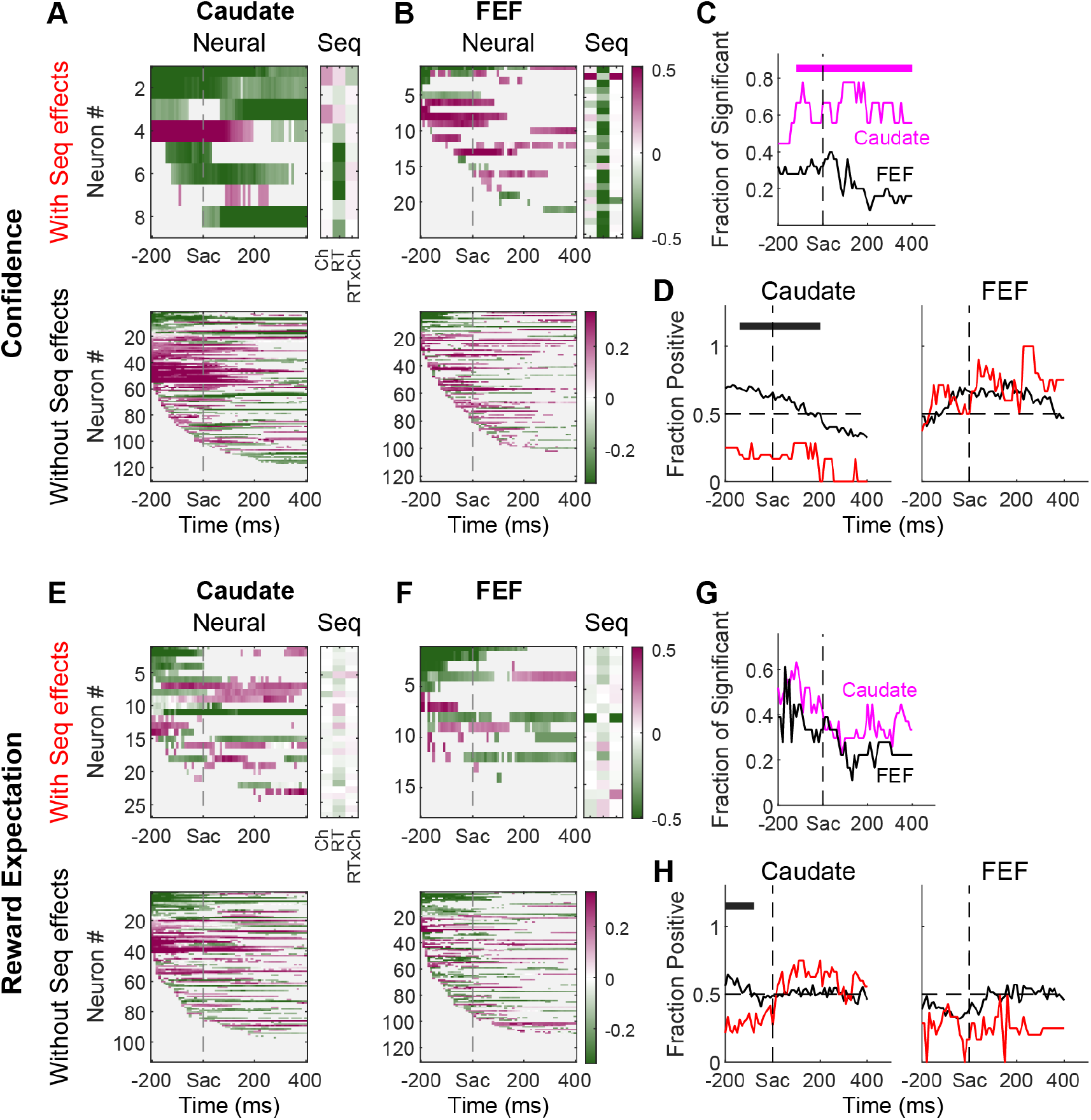
Relationship between observed sequential effects and neural modulation by choice confidence or reward expectation. **(A, B)** Left panels: partial correlation coefficients for choice confidence in caudate (A) and FEF (B) neurons recorded when there was (top) or was not (bottom) a confidence-dependent sequential effect. Right panels: regression coefficients for the sequential effects on choice, RT, and RT for the same choice. **(C)** Fractions of neurons showing significant partial correlation in sessions with choice confidence-dependent sequential effects. Horizontal bar indicates when there was a significantly higher fraction in caudate samples (Chi-Square test, *p* < 0.05). **(D)** Fractions of neurons showing positive partial correlation in sessions with (red) or without (black) choice confidence-dependent sequential effects. Horizontal bars indicate when the fraction was higher for the black curve (Chi-Square test, *p* < 0.05). **(E-H)** Same format as A-D. Neural data show partial correlation coefficients for reward expectation for sessions with (red) or without (black) reward expectation-dependent sequential effects.

In general, neural signatures of choice confidence and reward expectation were evident in both brain regions and in sessions both with and without behavioral sequential effects sensitive to those factors. Moreover, we did not detect a consistent relationship between the strength of neural representation (amplitude of the partial correlation coefficient) and the magnitude of sequential effects (regression coefficient from behavior) across these neurons, for either choice confidence or reward expectation. However, there was some evidence for a more reliable representation of behaviorally relevant evaluative signals in caudate than in FEF. First, confidence-related neural signals were more likely to be found in caudate than in FEF in sessions with confidence-related sequential behavioral effects (although note that there were fewer caudate than FEF neurons recorded in these sessions; Figure 8C). Second, in these sessions with confidence-dependent sequential choice effects, the caudate neurons were more likely to show negative partial correlation coefficients for choice confidence in most of the peri-saccade period, compared to when such a sequential effect was absent (Figure 8A and D). Third, in sessions with reward expectation-dependent sequential behavioral effects, the caudate neurons were more likely to show negative partial correlation coefficients for reward expectation in the pre-saccade period, compared to when such a sequential effect was absent (Figure 8E and H and Figure 8-Supplement 1). We did not detect any similar tendencies for FEF recordings. These results imply that evaluative signals represented in caudate activity might be more directly related to the monkeys’ online adjustments than those represented in FEF activity.

## Discussion

Choice confidence and reward expectation are both important quantities for post-decision evaluation, but they are often perfectly correlated in many common decision tasks. Using a task design with manipulations of sensory uncertainty and reward sizes, we were able to partially decorrelate and therefore identify distinguishable representations of these two conceptually distinct quantities. Focusing on post-decision activity in previously recorded FEF and caudate neurons (Doi et al., 2020; Fan et al., 2020), we observed that: 1) choice confidence and reward expectation were represented in both brain regions; 2) these representations were present around saccade onset in neurons that did and did not encode choice; 3) these representations were more prevalent in caudate than FEF neurons, especially for choice confidence; 4) the dominant sign of neural modulation by choice confidence followed different time courses in FEF and caudate neurons, especially for neurons also encoding choice; and 5) the evaluative signals in caudate neurons were more tightly linked to the monkeys’ decision adjustments in consecutive trials. These results provide new perspectives on previously reported cognitive signals in post-decision FEF and caudate activity patterns and further demonstrate functional differences between these two regions in decision evaluation.

Previous studies have shown that post-decision FEF and caudate neural activity is sensitive to various cognitive signals, including choice value (Kawagoe et al., 1998; Lau and Glimcher, 2008; Seo et al., 2012), task difficulty (Ding and Gold, 2010, 2012a; Teichert et al., 2014), confidence (Middlebrooks and Sommer, 2012; Yanike and Ferrera, 2014a), and accuracy-related risk (Yanike and Ferrera, 2014b). A simple hypothesis is that these different signals reflect the same underlying computations but are expressed differently under different task contexts. Our results, using a single task design, argue against this simple hypothesis by demonstrating that neural representations of at least two conceptually distinct signals co-exist in two brain regions that are well known to be involved in decision-making. Extrapolating from these results, it seems likely that even more diverse types of evaluative signals are present in the decision network, which includes other cortical areas, midbrain dopamine neurons, and superior colliculus (Kepecs et al., 2008b; Kiani and Shadlen, 2009b; So and Stuphorn, 2016; Lak et al., 2017, 2020b, 2020a; Odegaard et al., 2018). In principle, these diverse signals can be flexibly employed to adapt a decision-maker’s strategy to a wide range of decision goals. For example, the accuracy/confidence-related signals can be more readily used to evaluate performance when the goal is to maximize accuracy, detect a change in environments (Yu and Dayan, 2005; Nassar et al., 2012), implement multi-stage decisions (van den Berg et al., 2016; Desender et al., 2019a), or seek more information (Desender et al., 2019b). In contrast, reward expectation/risk-related signals can be more readily used when the goal is to maximize reward rate (Bogacz, 2007; Feng et al., 2009; Simen et al., 2009; Fan et al., 2018).

Alternatively, the presence of both choice confidence and reward expectation may be expected solely because choice confidence is a computational precursor of reward expectation. That is, FEF and caudate neurons are both involved in computing reward expectation from choice confidence, but only one of the two quantities may be functionally important. This interpretation is not consistent with our behavioral results of the monkeys’ sequential adjustments, which were sensitive to both confidence and reward expectation. However, because our datasets were obtained from monkeys trained extensively to base their decisions on the sensory and reward-context information presented for each trial, these sequential effects were relatively weak and occurred infrequently, thus limiting our ability to thoroughly test these alternative ideas. At an anecdotal level, our monkeys who showed sequential effects exhibited sensitive to choice confidence in some sessions, but reward expectation in others (Figure 3-Supplement 1). It will be interesting to track more systematically how a subject adjusts decision goals and tunes their decision strategy during learning, as well as if and how distributions of evaluative signals in the brain change accordingly.

Given the extensive projection from the FEF to the caudate, it is not surprising that the two regions share many functional similarities, particularly for decision-making. For example, we and others have shown previously that both the FEF and caudate carry information related to decision formation, such as uncertain sensory evidence (Kim and Shadlen, 1999; Ding and Gold, 2010, 2012a; Ding, 2015), values for potential outcomes (Kawagoe et al., 1998; Lauwereyns et al., 2002b, 2002a; Roesch and Olson, 2003; Samejima et al., 2005; Ding and Hikosaka, 2006; Lau and Glimcher, 2008), and the combination of them in complex decisions (Fan et al., 2020). The pre-decision activity in both regions is linked causally to decision behavior (Moore and Fallah, 2001; Ding and Gold, 2012b; Santacruz et al., 2017; Bollimunta et al., 2018; Doi et al., 2020). The similarity also extends to decision evaluation, as we show here that both regions carry information about choice confidence and reward expectation.

Despite these similarities, it is also clear that the caudate is not simply a relay station for FEF output. There are many notable regional differences even when the two regions are compared on the same task and in the same animals. For example, for a simple saccade task with reward manipulations, reward expectation-related information tends to be multiplexed with choice-selective activity in FEF, whereas it is encoded directly by the activity of a subset of caudate neurons (Ding and Hikosaka, 2006). FEF and caudate activity encoding reward context information also shows different temporal dynamics before a saccade can be planned (Ding, 2015). For a visual motion discrimination task, pre-decision FEF activity reflects motion evidence accumulation until a threshold level that is related to decision commitment, whereas caudate activity follows evidence accumulation only in the earlier phase of decision process (Ding and Gold, 2010, 2012a; Ding, 2015). For the asymmetric-reward motion discrimination task used here, FEF activity is more directly link to monkeys’ reward biases in evidence accumulation (Fan et al., 2020). Our new results document additional regional differences in decision evaluation. More specifically, the greater prevalence of choice confidence signals in caudate activity and the slightly tighter link between caudate activity and the monkeys’ sequential behavioral adjustments support the idea that the caudate is more directly involved in tuning the decision process based on evaluations of past decisions. This idea is further supported by previous observations that post-action caudate microstimulation can gradually bias RTs of a specific saccade (Nakamura and Hikosaka, 2006; Williams and Eskandar, 2006) and that caudate microstimulation during decision formation induces behavioral effects that mimics the monkeys’ voluntary reward bias strategies (Doi et al., 2020).

Another observation inconsistent with the relay hypothesis is the sign difference between FEF and caudate neurons, particularly for choice confidence representations (Figure 7D). Because the FEF-caudate projection is excitatory, the sign difference reflects either intermediary inhibition or another source of neural inputs to the caudate neurons. A potential candidate for the former are the striatal interneurons. Because these neurons are sparse relative to the striatal projection neurons that we recorded, future recordings using cell-type specific sampling techniques are needed to determine the involvement of striatal interneurons in decision-related computations. From outside the striatum, a potential source of choice confidence-related signals is the supplementary eye field, which has projection fields in the caudate that overlap with those of FEF and where neural activity is mostly negatively correlated with confidence on a value-based decision task (Parthasarathy et al., 1992; So and Stuphorn, 2016).

It should be noted that our results were based on our best estimates of choice confidence and reward expectation, which are internally generated quantities. Future studies with more direct measurements of choice confidence and reward expectation are needed to examine the neural representation of these evaluative signals in a more systematic manner. For example, our asymmetric reward design could be paired with post-decision wager that targets either choice confidence or reward expectation explicitly (Kiani and Shadlen, 2009b; Fetsch et al., 2014) or use a different task design that requires the subjects to report their relative choice confidence or reward expectation directly (Kiani et al., 2014b; Boldt and Yeung, 2015). Given the likely obstacles in training monkeys to perform these kinds of tasks, the advancement of intracranial recordings in human subjects may offer unprecedented opportunities to understand how decision evaluation is implemented in the brain.

In summary, we used a task design with independent manipulations of sensory evidence and reward associations to de-couple choice confidence and reward expectation. We found that a substantial fraction of neurons in the caudate nucleus and FEF encode these two different evaluative signals in their post-decision activity, but with regional differences in their prevalence, time course, and associations with behavior. These results highlight the diversity of signals and brain regions that contribute to how decisions are formed, evaluated, and adjusted to achieve particular goals.

## Materials and Methods

### Subjects, behavioral task, data acquisition, and DDM model fitting

The data sets for the present study were identical to those reported previously (Fan et al., 2020). The original report focused on neural activity during decision formation (i.e., after motion onset and before the saccadic response). The present study focused on neural activity around saccade onset that can encode evaluation of the decision. Details of subjects, the behavioral task, data acquisition, and DDM model fitting can be found in three previous reports (Fan et al., 2018, 2020; Doi et al., 2020) and are summarized here.

Briefly, a trial began with a central fixation point presentation (Figure 1a). Once the monkey acquired and maintained fixation, two choice targets were presented to indicate the two possible motion directions. After a random delay, the fixation point was dimmed and a random-dot kinematogram was shown (“motion onset”) with randomly interleaved motion direction and motion strength (coherence). The monkey reported the perceived motion direction by making a self-timed saccade to the corresponding choice target. Two asymmetric reward contexts were alternated in a block design. In Contra-LR blocks, the choice contralateral to the recording/stimulation site was paired with large reward (LR). In Ipsi-LR blocks, the choice ipsilateral to the recording/stimulation site was paired with large reward. The other choice was paired with small reward. The reward context for the current block was signaled to the monkey at the first trial. Three monkeys were extensively trained on this task. Single-unit recordings were obtained in the FEF and caudate nucleus (in separate sessions) while monkeys performed the task. DDM model fitting was performed, separately for each session, using the maximum *a posteriori* estimate method and prior distributions suitable for human and monkey subjects (Wiecki et al., 2013). The same fitting results were reported previously (Fan et al., 2018, 2020).

### Computation of choice confidence and reward expectation

Following previous literature (Kiani and Shadlen, 2009a), we defined choice confidence as the estimation of accuracy on average given the current choice and decision time, as following:

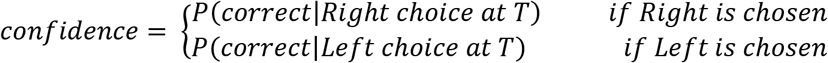

where *T* is the decision time that equals RT minus non-decision time (estimated from DDM fits). The righthand side was computed by marginalizing over all possible coherences. For example, for rightward choices:

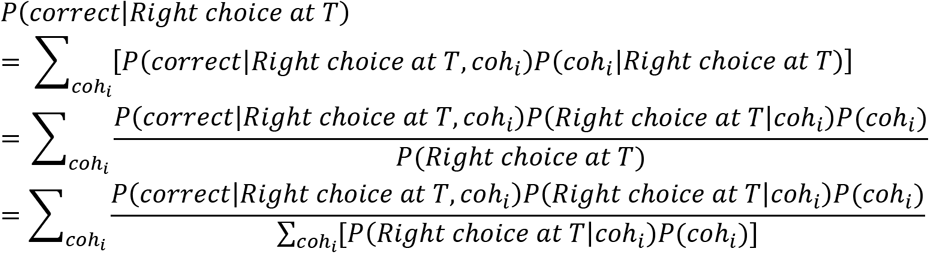

where *coh_i_* is signed coherence (+/- for rightward and leftward motion).

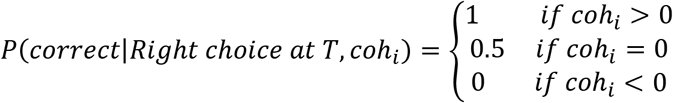

In our task design, each coherence had an equal chance of appearance, except that *coh*=0 happened twice as often as the other coherences:

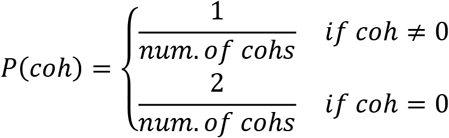

*P(Right choice at T*|*coh_i_*) was obtained by DDM simulation using the best-fitting parameters, as illustrated in Figure2-supplement 1. For each coherence, we obtained the probability of the decision variable (DV) attaining a value *x* at time *t, P*(*DV*(*t*) = *x*), using the best fitting DDM parameters of each session and reward context.

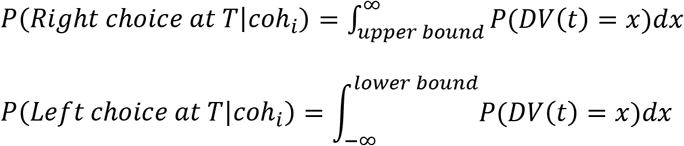

After obtaining an estimate of choice confidence,

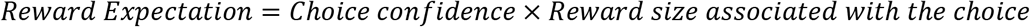

To standardize across sessions with different juice volumes, we normalized reward size by the volume of the smaller reward for each session. That is, for each session the small reward was assigned a reward size of 1 and the large reward was assigned a value equal to the large-small reward ratio.

### Measurement of sequential effects

Due to the limited number of error trials, we only used trials after correct responses. We focused on three types of sequential effects that depended on evaluative signals. First, the current choice may either repeat the previous choice or switch to the other choice. To assess this effect, we performed logistic regression using the following function:

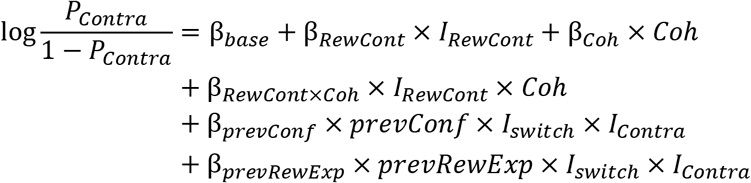
where *P_Contra_* is the probability of choosing the contralateral option; *Coh* is the signed coherence of current trials (+/- for motion towards contralateral/ipsilateral direction); and *prevConf* and *prevRewExp* are the choice confidence value and reward expectation value, respectively, in the previous trial.

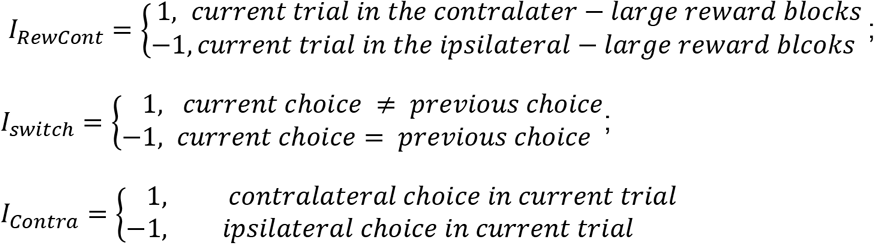

β_*prevConf*_ > 0 implies that high confidence in the previous trial increases the chance of switching to the opposite choice in the current trial. β_*prevRewExp*_ > 0 implies that high reward expectation in the previous trial increases the chance of switching to the opposite choice in the current trial. In Figure 3, we normalized the effect of choice confidence (β_*prevConf*_) and reward expectation (β_*prevRewExp*_) by motion sensitivity (β_*Coh*_), so that the sequential effects on choice are in the unit of 1/coh.

Second, the current choice may be sped up or slowed down depending on evaluation of the previous trial. Third, the speed-up/slow-down may also depend on a combination of evaluation and whether to switch choices. To assess these effects on RT, we performed the following linear regression:

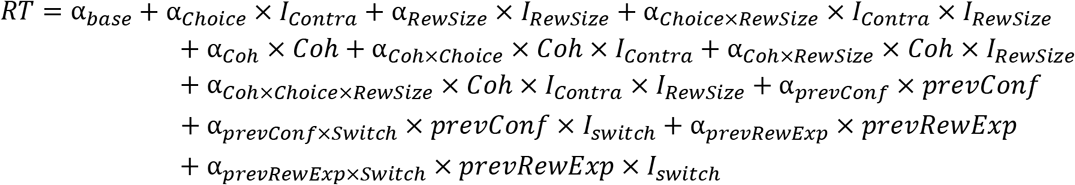
where *Coh* is the un-signed motion coherence in the current trials (positive for both directions); *prevConf* and *prevRewExp* are the confidence value and reward expectation value in the previous trial.

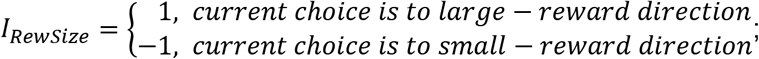

α_*prevConf*_ > 0 means that high confidence in the previous trial increases (a.k.a., slows down) the RT in the current trial, regardless of the saccade direction. α_*prevConf×Switch*_ > 0 means that high confidence in the previous trial increases (slows down) the RT in the opposite direction while decreases (speeds up) RT in the same direction in the current trial. Similar interpretation with α_*prevRewExp*_ and α_*prevRewExp×Switch*_.

Because choice confidence and reward expectation can still be correlated, even after the decoupling, we used a model-selection procedure to infer their individual influences. For each session, we fitted both choice and RT data to four sets of models. The first set (“Both”) included the logistic and linear regressions described above and assumed that both choice confidence and reward expectation were involved. The second and third sets (“ConfOnly” and “RewExpOnly”) were reduced versions of the first set and assumed that either choice confidence or reward expectation were involved, respectively, but not both. The last set (“Neither”) further reduced the regressions by assuming that neither choice confidence nor reward expectation was involved. For each session, we deemed the fitted values for the set with the smallest Akaike Information Criterion value as the best estimates of sequential effects.

We considered that a session showed choice confidence-dependent sequential effects if any confidence-related regression coefficient was non-zero (p<0.05, t-test). Similarly, we considered that a session showed reward expectation-dependent sequential effects if any reward expectation-related regression coefficient was non-zero. We used these lenient criteria to look for any trends of sequential effects in these extensively trained monkeys. These criteria also meant that a session considered to show sequential effect did not necessarily show non-zero coefficients for all relevant regressors.

### Neural data analysis

We focused on neural activity between 200 ms before saccade onset (i.e., near decision commitment) to 400 ms after saccade onset. For each neuron, we measured the average firing rates in 300-ms time windows with 10-ms steps. For each time window, we performed two partial (Spearman) correlations: 1) between firing rates and choice confidence while removing the effect of reward expectation, and 2) between firing rates and reward expectation while removing the effect of choice confidence. Significance was assessed at *p*=0.05. Chi-square tests were performed to compare fractions of significant modulation at each time window between conditions, with corrections for multiple comparisons. Figures 6 and 7 show the results based on data from correct trials only. Similar results were obtained from all trials (not shown).

We tested the effects of two potential confounds. First, because choice confidence and reward expectation are both affected by reward biases, it is possible that reward context modulation alone may cause measurable correlations between firing rate and choice confidence or reward expectation. To minimize such a possibility, we imposed an additional criterion that modulation by choice confidence or reward expectation must be accompanied by modulation by decision time. For each time window and choice, we computed the correlation between firing rates and decision time for the two reward contexts separately and jointly. We considered a significant modulation by decision time to be present if any of the three correlation coefficients were non-zero (*p*<0.05).

Second, we assessed whether a subjective reward ratio, different from the actual ratio of juice volume, may provide a more accurate measurement of reward expectation and significantly affect the prevalence of reward expectation modulation of neural activity.

We computed new reward expectation with reward ratio ranging from 1 to 3.5 and operationally defined the “best” reward ratio as the value associated with the largest correlation between firing rate and reward expectation (Figure 7-Supplement 2A). For this analysis, we used three representative epochs: 1) a pre-saccade 100ms window beginning at 100ms before saccade onset, 2) a peri-saccade 300 ms window beginning at 100 ms before saccade onset, and 3) a post-saccade 400 ms window beginning at saccade onset (all the epochs end before reward delivery).

## Acknowledgements

We thank Jean Zweigle for animal care. This work was supported by NIH National Eye Institute (R01-EY022411; L.D. and J.I.G), University of Pennsylvania (University Research Foundation Pilot Award; L.D.), and Hearst Foundations Graduate student fellowship (Y.F.).

---

**Figure 2-supplement1.**
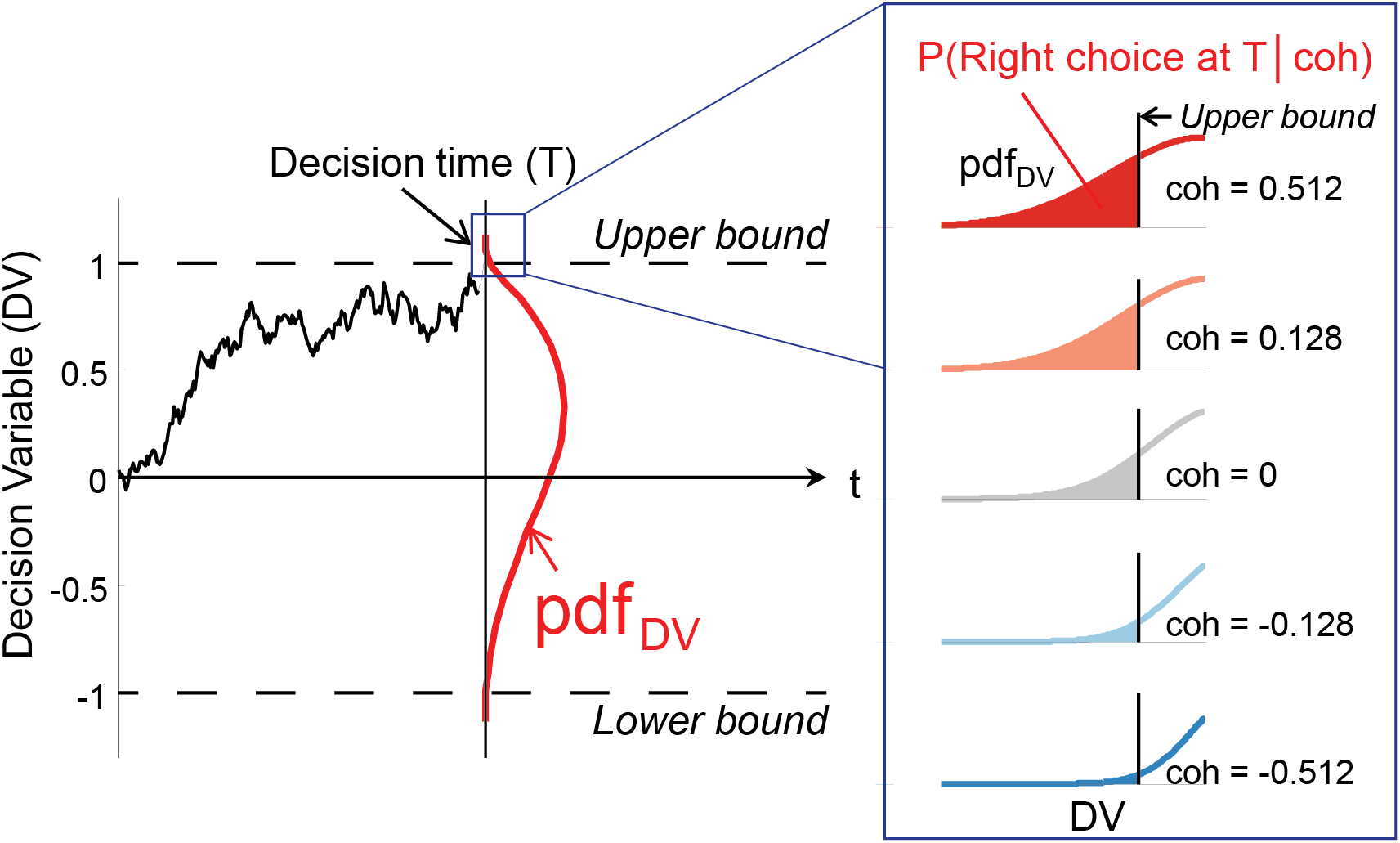
Related to “Computation of choice confidence and reward expectation” in Methods: Computing the probability of making a rightward choice at time T for a given motion coherence. Schematic illustrating how to compute the probability of making a rightward choice at time *T* for a given motion coherence *(P(Right* choice at *T* | *coh))* using the DDM framework. For each coherence, obtain the probability of decision variable (DV) attaining value x at time t (red curve), then compute the area underneath the probability function when DV > *Upper* bound (shaded area). For leftward choices, compute the area underneath the probability function when DV < *Lower* bound.

**Figure 3-supplement 1.**
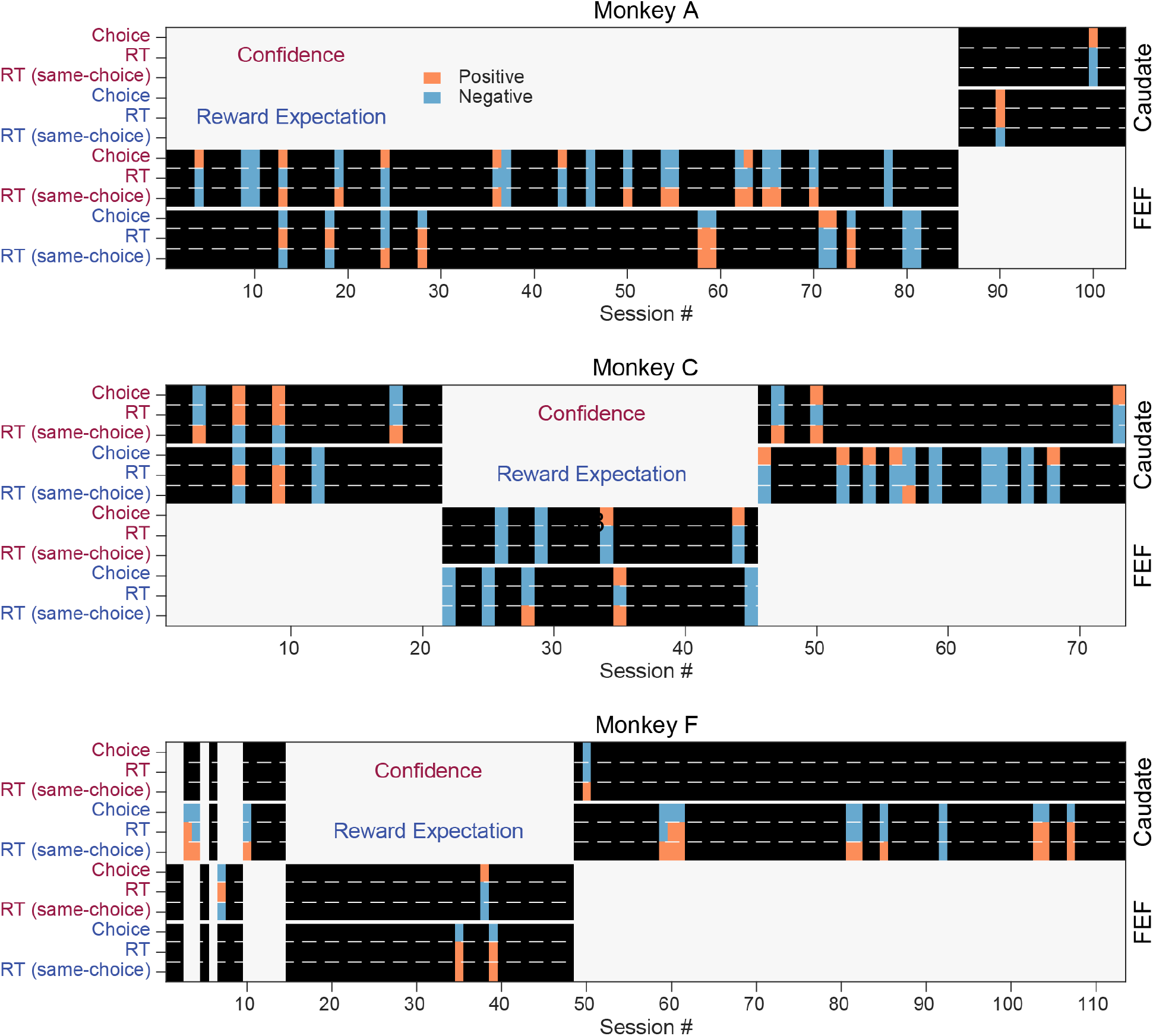
Sequential behavioral effects identified for each session (abscissa) and monkey (panel). For each panel, the top and bottom halves show caudate and FEF recording sessions, respectively. For each session, the signs of the regression coefficients are shown as blue (negative) or orange (positive). For sessions that were best described by “ConfOnly” or “RewExpOnly” models, the coefficients for the other evaluative signal were set as zero (black). For sessions that were best described by the “Neither” model, all coefficients were set as zero.

**Figure 6-Supplement 1.**
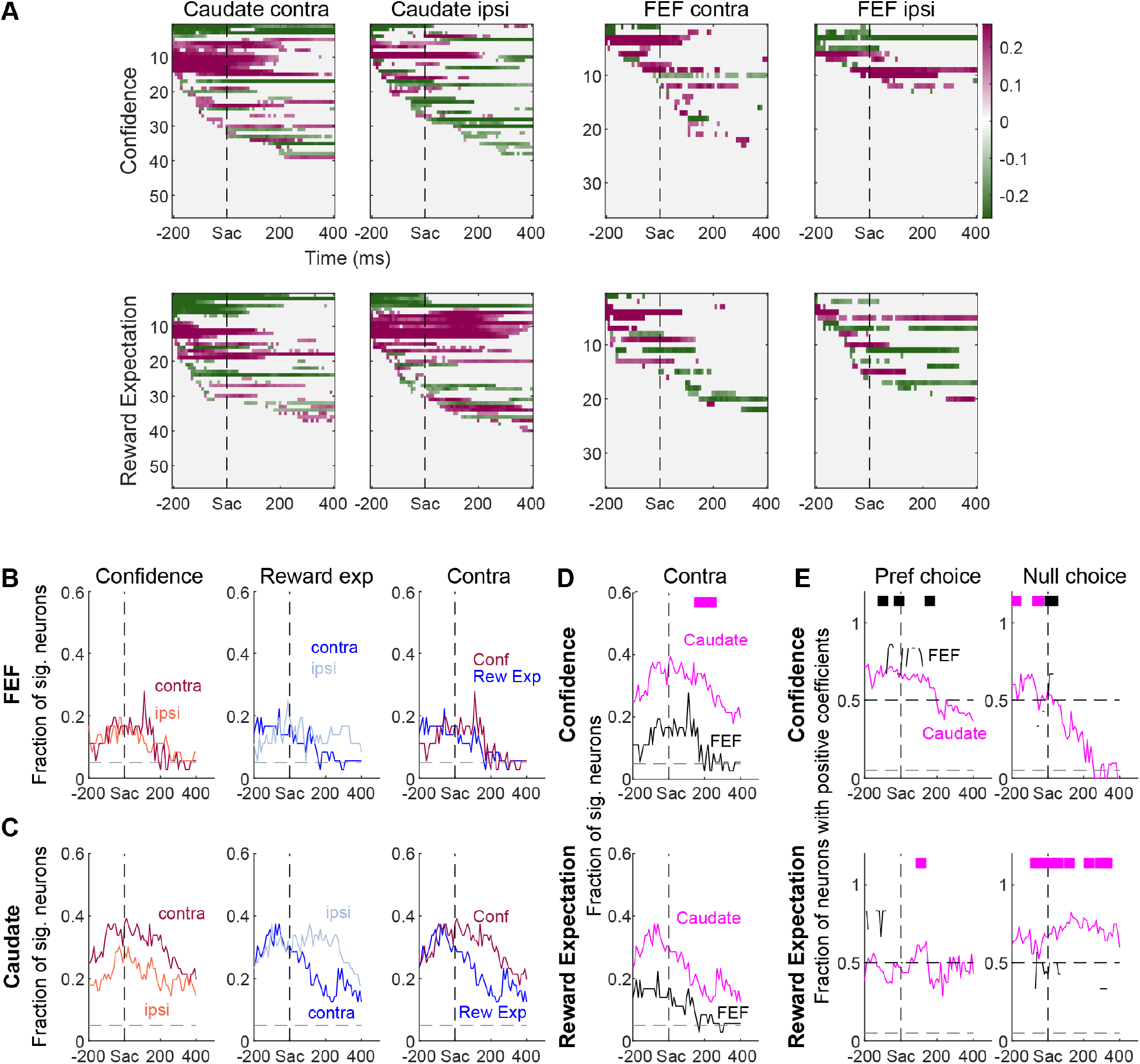
Partial correlation results for neurons without choice-selective activity around saccade onset. **(A)** Heatmap of correlation coefficients. Same format as Figure 6. **(B-E)** Comparisons of fractions. Same format as Figure 7. Note that, in many time bins, there were too few FEF non-choice-selective neurons that showed significant non-zero coefficients for the grouping by signs.

**Figure 7-Supplement 1.**
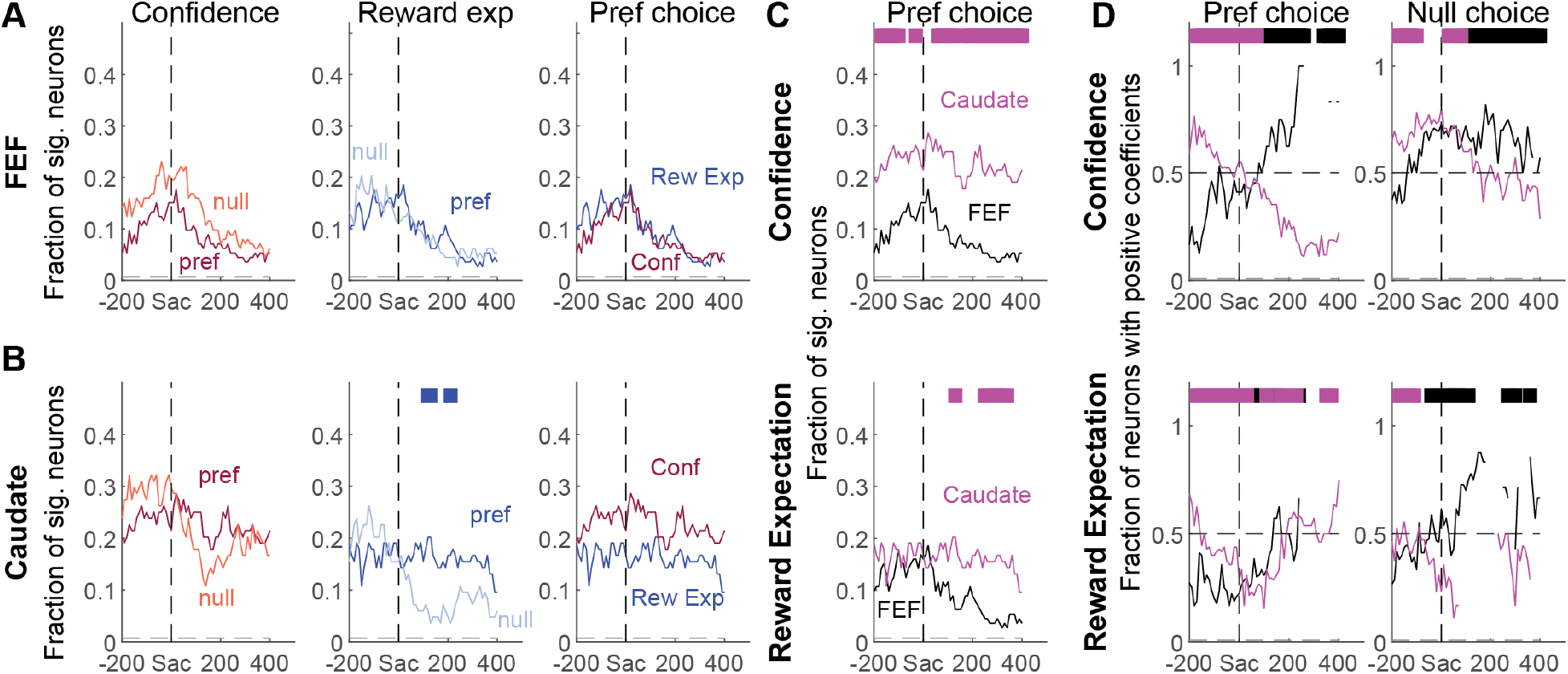
Partial correlation results for neurons with choice-selective activity around saccade onset, with an additional criterion of modulation by decision time. Same format as Figure 7.

**Figure 7-Supplement 2.**
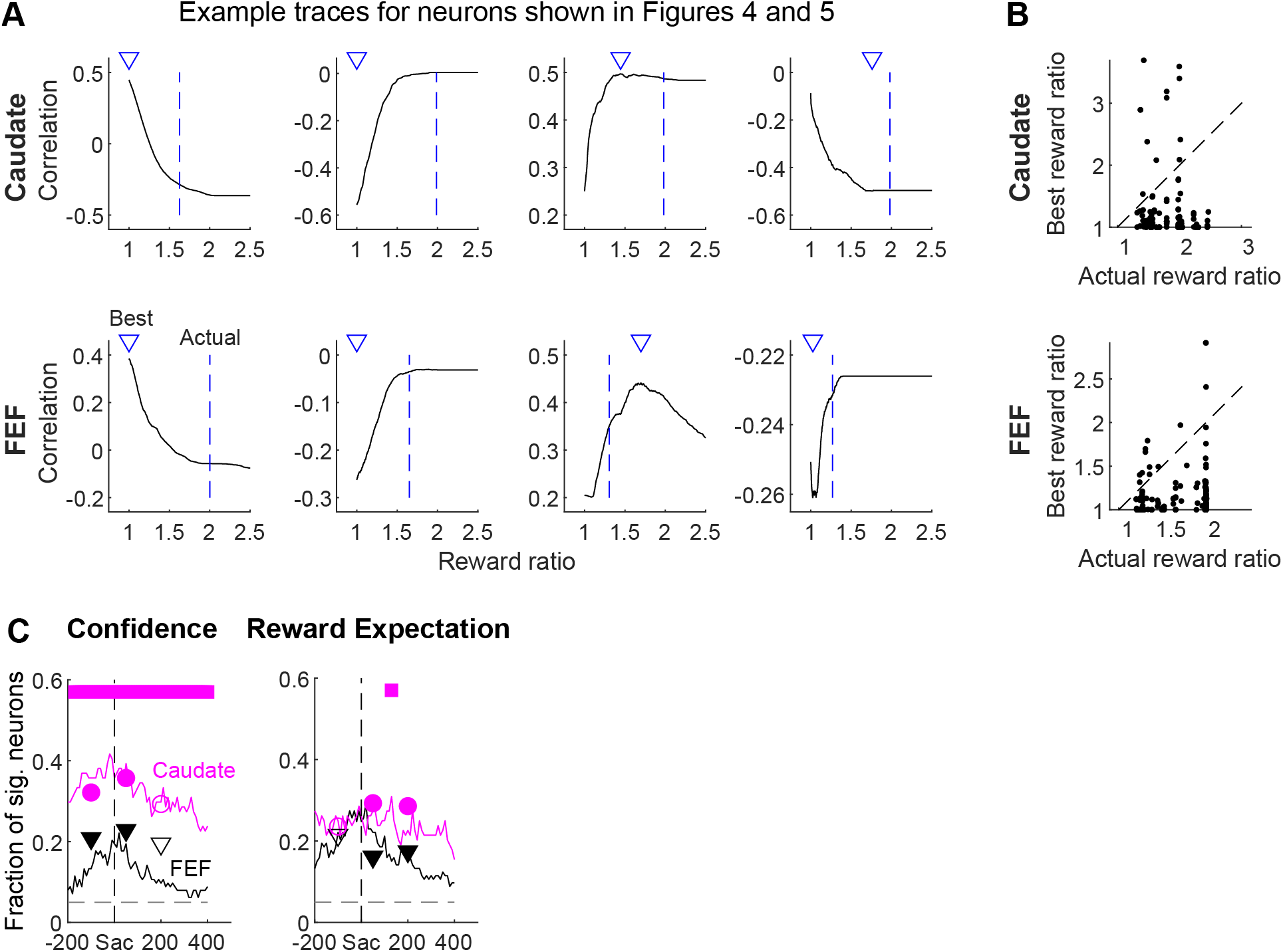
The regional differences were not due to estimation errors for the subjective reward ratio. **A,** Illustration of the identification of the best reward ratio (triangles) in the correlation function between firing rates and reward expectation values calculated with different reward ratios. Dashed lines: the actual ratio in juice volume. **B,** Scatterplots of the best and actual reward ratios estimated using firing rates in presaccade, peri-saccade, and post-saccade epochs for all sessions. Note that the best reward ratio is expected to be near one for activity modulated only by choice confidence. **C,** Comparisons between the original fraction traces (same as in Figure 7C) and the fractions measured using best reward ratios (circles: caudate samples, triangles: FEF samples) for the three epochs. Filled symbols: significant regional difference (Chisquare test, p<0.05). Note that the same patterns of regional difference remained.

**Figure 8-Supplement 1.**
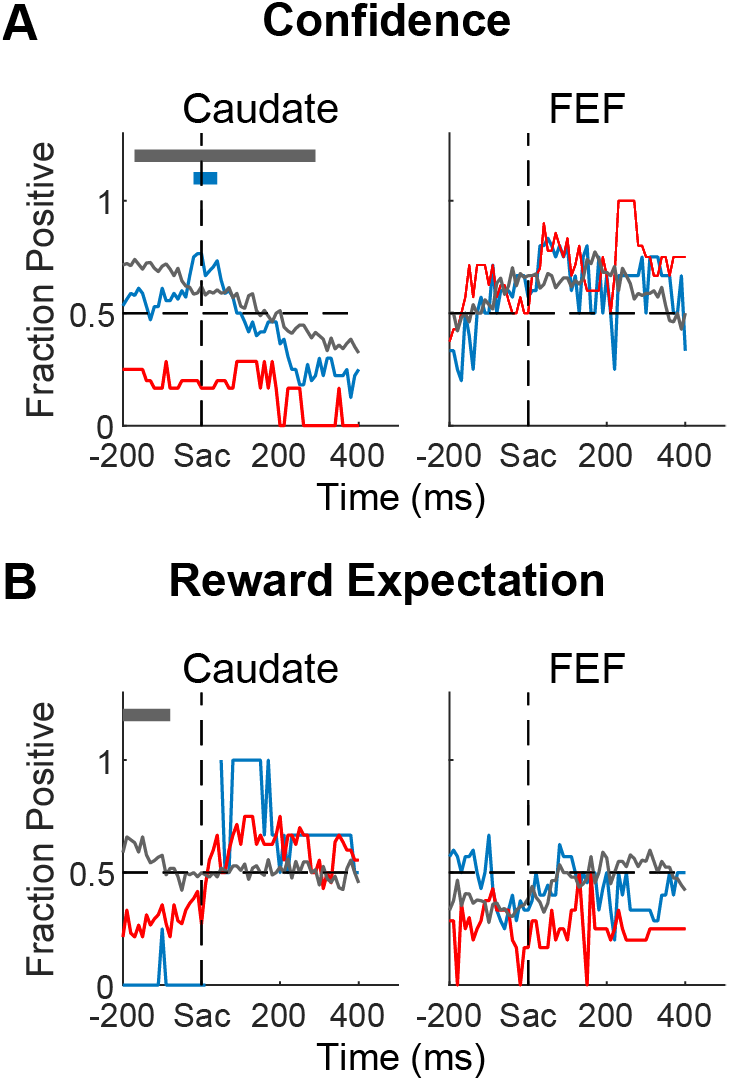
Additional comparisons of the signs of partial correlation coefficients between neural activity and confidence (A) or reward expectation (B) for sessions with different sequential effects. **A,** Same format as Figure 8D. Red: sessions with confidence-dependent sequential effects. Gray: sessions without any sequential effects. Teal: sessions with only reward expectation-dependent sequential effects. Horizontal bars indicate the time bins when the gray or teal curves were significantly higher (Chi-square test, *p* < 0.05). **B,** Same format as Figure 8H. Red: sessions with reward expectation-dependent sequential effects. Gray: sessions without any sequential effects. Teal: sessions with only confidence-dependent sequential effects. Horizontal bars indicate the time bins when the gray or teal curves were significantly higher (Chi-square test, *p* < 0.05).

